# Real-time cryo-EM data pre-processing with *Warp*

**DOI:** 10.1101/338558

**Authors:** Dimitry Tegunov, Patrick Cramer

**Author notes:** Correspondence should be addressed to D.T. and P.C.

## Abstract

The acquisition of cryo-electron microscopy (cryo-EM) data from biological specimens is currently largely uncoupled from subsequent data evaluation, correction and processing. Therefore, the acquisition strategy is difficult to optimize during data collection, often leading to suboptimal microscope usage and disappointing results. Here we provide *Warp*, a software for real-time evaluation, correction, and processing of cryo-EM data during their acquisition. *Warp* evaluates and monitors key parameters for each recorded micrograph or tomographic tilt series in real time. *Warp* also rapidly corrects micrographs for global and local motion, and estimates the local defocus with the use of novel algorithms. The software further includes a deep learning-based particle picking algorithm that rivals human accuracy to make the pre-processing pipeline truly automated. The output from *Warp* can be directly fed into established tools for particle classification and 3D image reconstruction. In a benchmarking study we show that *Warp* automatically processed a published cryo-EM data set for influenza virus hemagglutinin, leading to an improvement of the nominal resolution from 3.9 Å to 3.2 Å. *Warp* is easy to install, computationally inexpensive, and has an intuitive and streamlined user interface.

## INTRODUCTION

Modern cryo–electron microscopy (cryo–EM) allows structures of protein assemblies to be solved at an ever-increasing speed. Automation of the data acquisition process and improved instrument stability have reduced the time required to collect hundreds of thousands of particle images to mere days. As the field attracts more scientists and the research projects become more ambitious, data collection time on a high-end microscope will remain a scarce and expensive resource. To make the best use of microscope time, the operator needs to continuously monitor data quality. Because many artifacts only become evident in the later stages of processing, the results of as many processing steps as possible should be available quickly to enable informed decisions during data collection.

The standard cryo–EM processing pipeline begins with the alignment of dose-fractionated image frames to cancel out sample motion. This is followed by an estimation of the contrast transfer function (CTF). Particles are then selected and extracted from the images. These steps are usually referred to as ‘data pre-processing’. Subsequent 2D and 3D classification of the particles serves to separate different complex conformations. Finally, 3D refinement is performed to obtain a three-dimensional map that is used to build a model.

Separate tools exist that can each handle one of the pre-processing steps^1-4^. Software packages like Appion^5^, FOCUS^6^ and Scipion^7^ are built around these tools to provide a common interface. Several tools can also be combined, but the resulting toolchains are not user-friendly and are not performing at high speed. No toolchain today can reliably perform pre-processing of cryo–EM raw data at the speed of data collection. Only for exceptionally favorable samples data processing has been reduced to “one push of a button” by fully automated procedures^8, 9^. In the vast majority of cases, however, human expertise, creativity, and intervention remain required.

Here we present *Warp*, a new software that takes over all pre-processing steps for 2D and tomographic cryo–EM data. *Warp* offers improved, real-time algorithms and a fully visual, intuitive interface that assists the user with immediate feedback and an augmenting presentation during data collection and pre-processing. The new algorithms can handle both standard data collection strategies and advanced scenarios such as tilted 2D data collection^10^ or dose-symmetric tilt series^11^, in a transparent, unsupervised manner. Coupled with automated acquisition software^12, 13^, Warp provides a continuous low-latency stream of reliably picked and corrected particle images that can be seamlessly fed into 2D classification, *ab initio* reconstruction, and 3D refinement using other packages^9^. *Warp* offers an accurate impression of the data quality during data collection, monitors the microscope behavior, and can enable advanced data analysis before data acquisition is completed. In favorable cases, this can result in high-resolution cryo-EM structure solution during ongoing data collection.

## RESULTS

### Rationale of *Warp*

We aimed at providing a software package that enables the electron microscopy user to evaluate, correct, and process cryo-EM raw data immediately during data acquisition. The rationale was to provide a single, streaming interface between the data acquisition software that produces the raw data, and the existing software solutions for 2D classification and 3D refinement of pre-processed cryo-EM single-particle data, such as cryo-SPARC^9^ or RELION^8^. We called the resulting software package ‘*Warp*’ in reference to its highly efficient correction of object distortions that occur in cryo-EM and its GPU-based implementation that results in almost instantaneous output. Our rationale was that *Warp* should be used for the online evaluation, correction and processing of both 2D and tilt series cryo-EM data. *Warp* can be installed on standard platforms and operated by non-expert users via a streamlined user interface (UI) that has been developed in parallel to the underlying algorithms to augment their operation. *Warp* was designed to be widely applicable for biological data acquisition at any cryo-EM facility and substantially speeds up the process of cryo-EM structure determination with improved results.

### Overall design

A schematic of the computational steps carried out by Warp is provided in Fig. 1. For simplicity, we first describe the workflow for 2D data, before we describe the application to tomographic tilt series at the end of the results section. At the beginning of the pipeline, *Warp* reads any new data saved by the acquisition software. *Warp* then estimates and corrects the motion captured in the frames both globally and locally. Next, *Warp* fits a spatially resolved CTF model, enabling the assignment of local defocus values to any particles extracted from the micrograph later. *Warp* then uses a neural network-based approach to automatically pick particles from the corrected micrographs with very high accuracy. Finally, *Warp* exports the resulting dose-weighted particle images to a downstream structure determination program such as cryoSPARC^9^ or RELION^8^ that carry out 2D and 3D classification, map refinement and reconstruction. During pre-processing, *Warp* provides a comprehensive overview of all important data parameters, allowing the operator to tune the acquisition settings to achieve optimal results faster. In the following we will describe the most important components in more detail.

**Figure 1.**
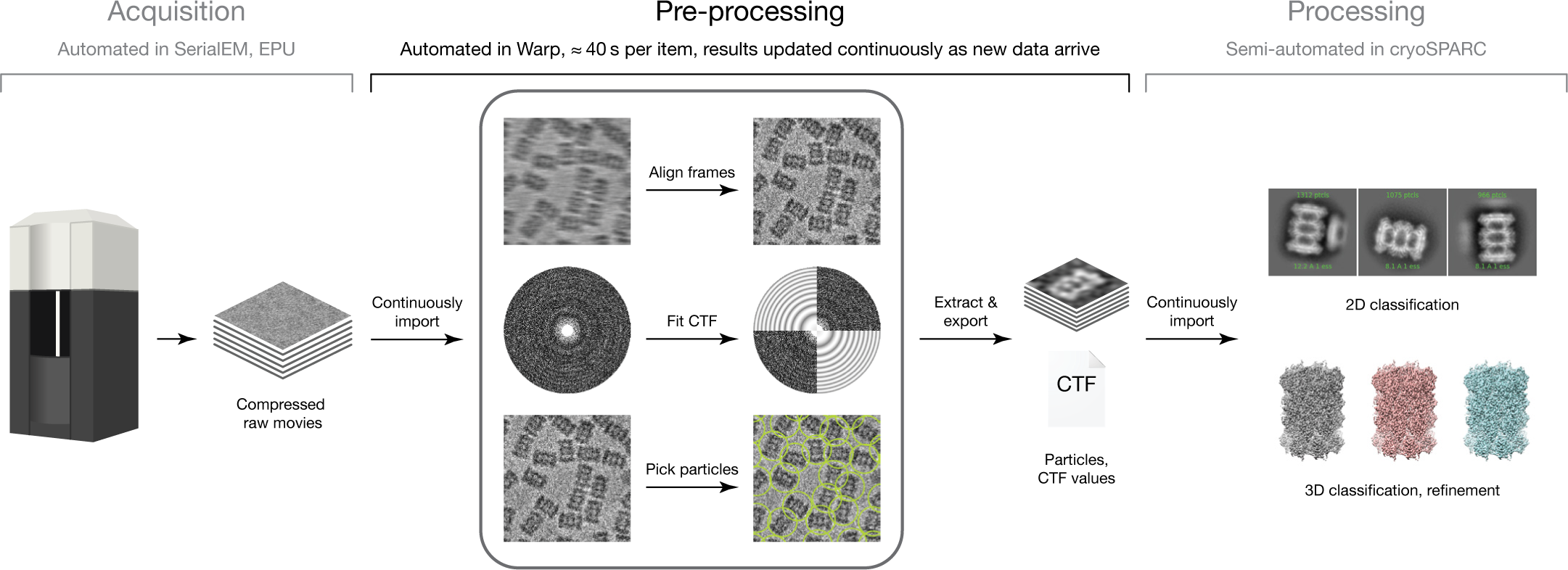
Warp handles all pre-processing steps to close a gap in the 2D cryo–EM pipeline. Data is acquired by the microscope in an automated fashion and stored as compressed stacks of movie frames. *Warp* continuously monitors its input folder for new files, and subjects them to all steps of the pre-processing pipeline: frame alignment, CTF estimation and particle picking. *Warp* writes out a stack of particles for each pre-processed micrograph and maintains a dynamically updated STAR file with references to all particles and their local CTF parameters. This file can be used as a data source in a tool such as cryoSPARC, which will periodically run subsequent processing steps like 2D classification and ab initio reconstruction on the latest set of particles.

### User interface

*Warp*’s UI is designed to help the user to comprehend and interact with the thousands of data objects generated routinely during cryo-EM data collection (Fig. 2). The ‘Overview’ tab displays various properties, such as defocus, estimated resolution, amount of motion, or particle count, for all processed micrographs or tilt series as interactive plots. The user can immediately grasp the statistical distribution, observe intrinsic patterns, and make an informed decision to manually adjust the acquisition parameters, e.g. to tune the lens astigmatism, increase the stage settling time, or skip a bad grid square. A filter range can be specified for every plotted parameter to automatically exclude lower-quality images from downstream processing. Any data point can be quickly inspected in more detail in a tab called ‘Fourier & Real Space’. Here, a display of the power spectrum and the CTF fit allows optimization of CTF fitting parameters. In the real-space view, a deconvolution filter (Methods, Fig. 7) can be instantly applied to micrographs to improve the contrast and make the particles more visible to the human eye. The defocus variation obtained through local CTF estimation can be overlaid semi-transparently. Particle picking results can be assessed in the context of a micrograph, and edited manually. Dedicated dialogs assist the user with tasks like micrograph list export, particle extraction, template matching, tomogram reconstruction, and neural network training.

**Figure 2.**
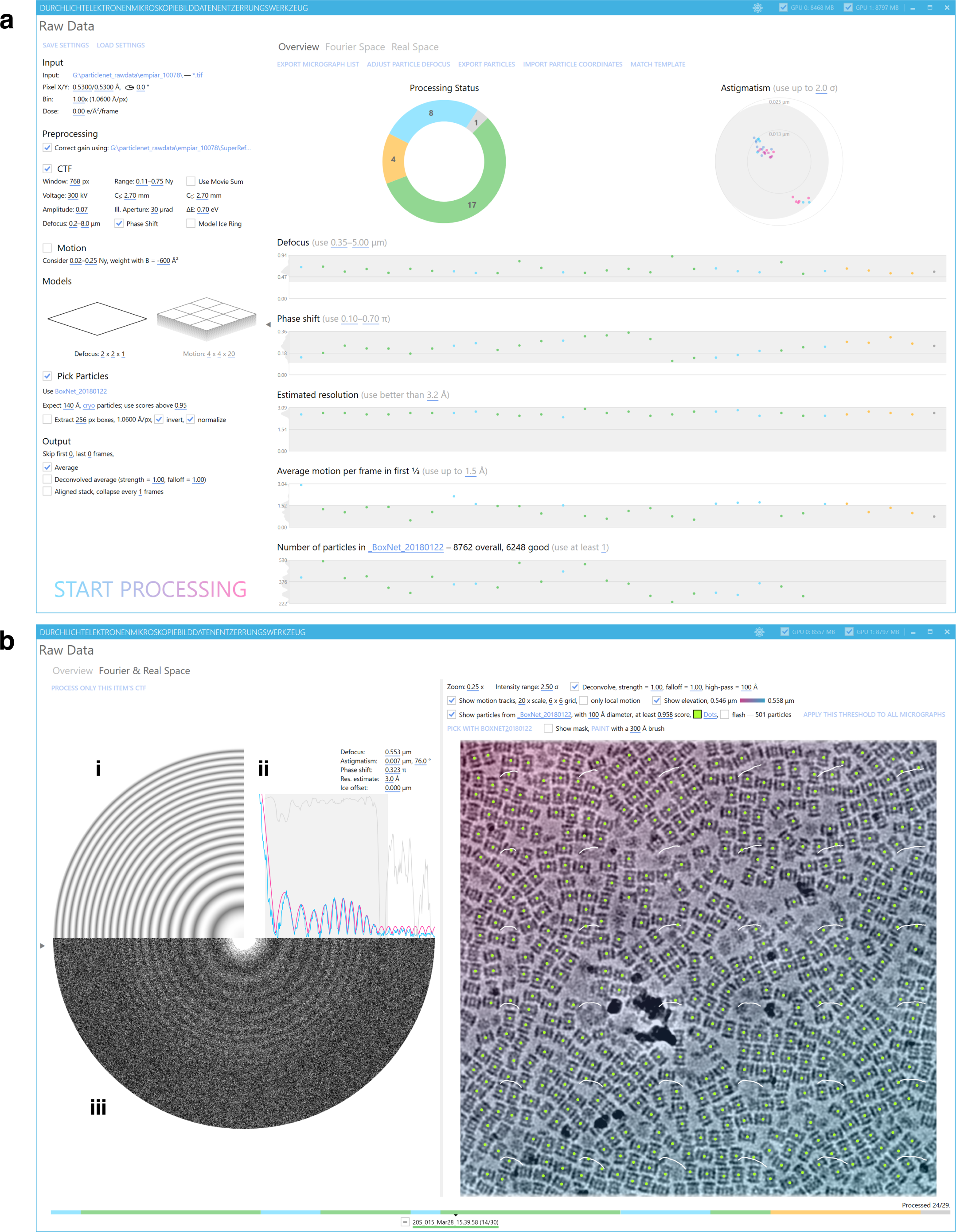
User interface of *Warp*. (a) The processing settings (left) specify all steps and parameters for online data evaluation, correction and processing. The ‘Overview’ tab (right) presents all important processing results and lets the user specify selection filters to remove low-quality data. (b) View of a single micrograph. In Fourier space (left), the simulated 2D CTF (i), the 1D power spectrum (PS) and its fit (ii), and the 2D PS (iii) are presented. The real space view (right) shows the aligned movie average with particle positions (green dots), motion tracks (white curves) and the defocus variation (transparent magenta-cyan overlay), and applies a deconvolution filter. Individual display elements can be shown or hidden. The navigation bar (bottom) shows the processing status for all items and allows to quickly switch between them as well as to manually exclude single items from processing.

### Motion correction

*Warp* generally represents space- and time-dependent parameters as coarse, uniform grids, on which a computationally cheap B-spline interpolation can retrieve any intermediate value (Methods). The motion, i.e. the translational shift observed between frames, is due to two effects: movement of the mechanical sample stage, and beam-induced motion (BIM). Stage movement causes global shifts over the entire field of view, whereas BIM leads to shifts between adjacent micrograph patches^14, 15^. While stage drift can lead to rapid shift changes between frames, BIM occurs more slowly after rapid relaxation during initial exposure^16^. *Warp* corrects for both global drift and local BIM at variable temporal resolution (Fig. 3). The strategy is similar to the one used by MotionCor2^1^, except that *Warp* does not apply any additional *a priori* assumptions about BIM beyond those imposed by the parameter grid resolution. As a result, *Warp* corrects very efficiently and thoroughly for the two types of motion that occur during cryo-EM data acquisition in any kind of support film hole morphology and orientation.

**Figure 3.**
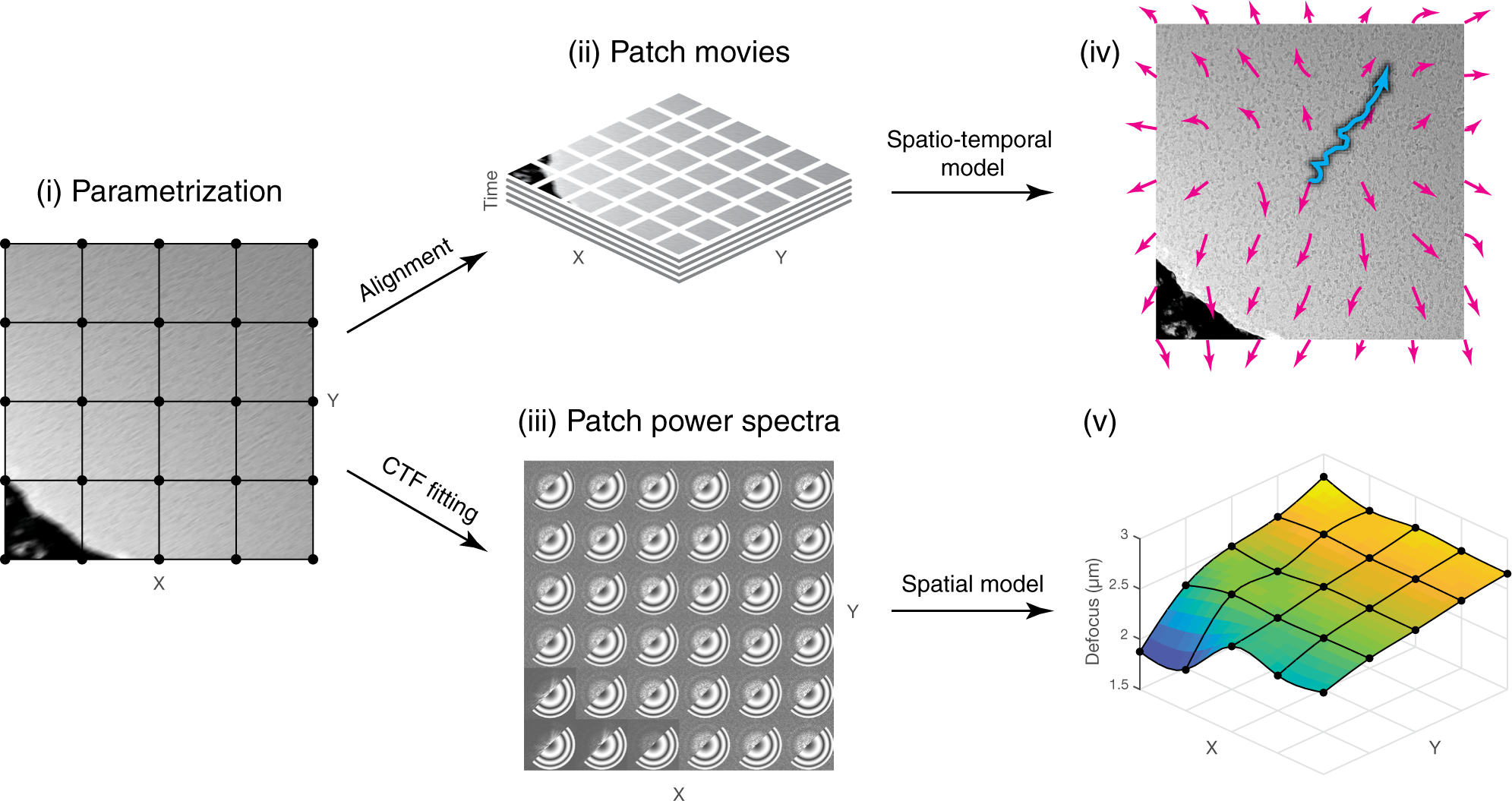
Motion and CTF model fitting by *Warp*. The unaligned, defocused movie (i) is parametrized with a coarse grid (black dots), divided into patches for the alignment (ii), and power spectra of these patches are computed (iii) for CTF fitting. The motion model (iv) includes 2 components: global motion (cyan trajectory) with fine temporal and no spatial resolution, and local motion (magenta trajectories) with coarse temporal, and fine spatial resolution. Both components are optimized to minimize the squared difference between the individual patch frames and their aligned average. The spatially resolved CTF model (v) is optimized to minimize the squared difference between the power spectra (iii, upper left part of each patch) and the simulated local 2D CTF (iii, bottom right part of each patch). Here, the defocus gradient follows the 40° tilt of the specimen, with the notable exception of the hole edge in the bottom left corner.

### Estimation of local defocus and resolution

The CTF model can be estimated based on the power spectrum (PS) of a micrograph. However, the defocus varies over the micrograph area due to stage inclination, uneven sample surface, or an uneven particle distribution along the optical axis^17^. *Warp* provides a flexible way to model local defocus variation in spatial and temporal dimensions without the need for *a priori* knowledge of particle positions. Instead of one global estimate, a tilted plane or a more complex geometry is fitted to the PS of a movie patch to estimate local defocus. A 1D average of all local power spectra rescaled to a common defocus value allows the user to easily assess whether fitting the more complex geometry recovered more Thon rings beyond the spatial frequencies used for the fitting. Thus, *Warp* goes beyond state-of-the-art CTF estimation by providing a spatially resolved model without the need for *a priori* knowledge of particle positions, and costly hyper-parameter tuning. The spatially resolved CTF model can converge on the correct solution for tilts as high as 60°, based on the inclination of the estimated defocus gradient. This makes *Warp* a useful tool for tilted 2D data collection, which has been shown to increase the resolution isotropy for samples with preferred orientation^10^. The useful resolution range of a micrograph is estimated as the spatial frequency where the fit quality falls below a threshold (Methods).

### Particle picking with BoxNet

The next step in cryo-EM structure determination is the accurate selection of single particles from the corrected micrographs. *Warp* includes a novel particle picking routine based on a machine learning algorithm (Methods). For several years, the computer vision community has been using convolutional neural nets (ConvNets) to vastly outperform template matching in object recognition tasks^18, 19^. First attempts to apply ConvNets to the particle picking problem in cryo-EM have shown performance on par with traditional template matching approaches^20^. Today, deep residual network (ResNet) architectures enable the training of arbitrarily deep models^21^. *Warp* employs ‘BoxNet’ – a fully convolutional ResNet architecture with 72 layers, implemented in TensorFlow 1.5^22^. BoxNet was trained with data from the EMPIAR raw data repository^23^ and synthetic data simulated from PDB^24^ structures with a molecular weight range of 0.064–18 MDa. As a result of these efforts, the pre-trained neural network bundled with *Warp* performs well on many new particle species and is able to accurately mask out high-contrast artifacts, such as ethane. The performance of BoxNet compares favorably with available tools when representative single-particle cryo-EM data are used as input (Fig. 4).

**Figure 4.**
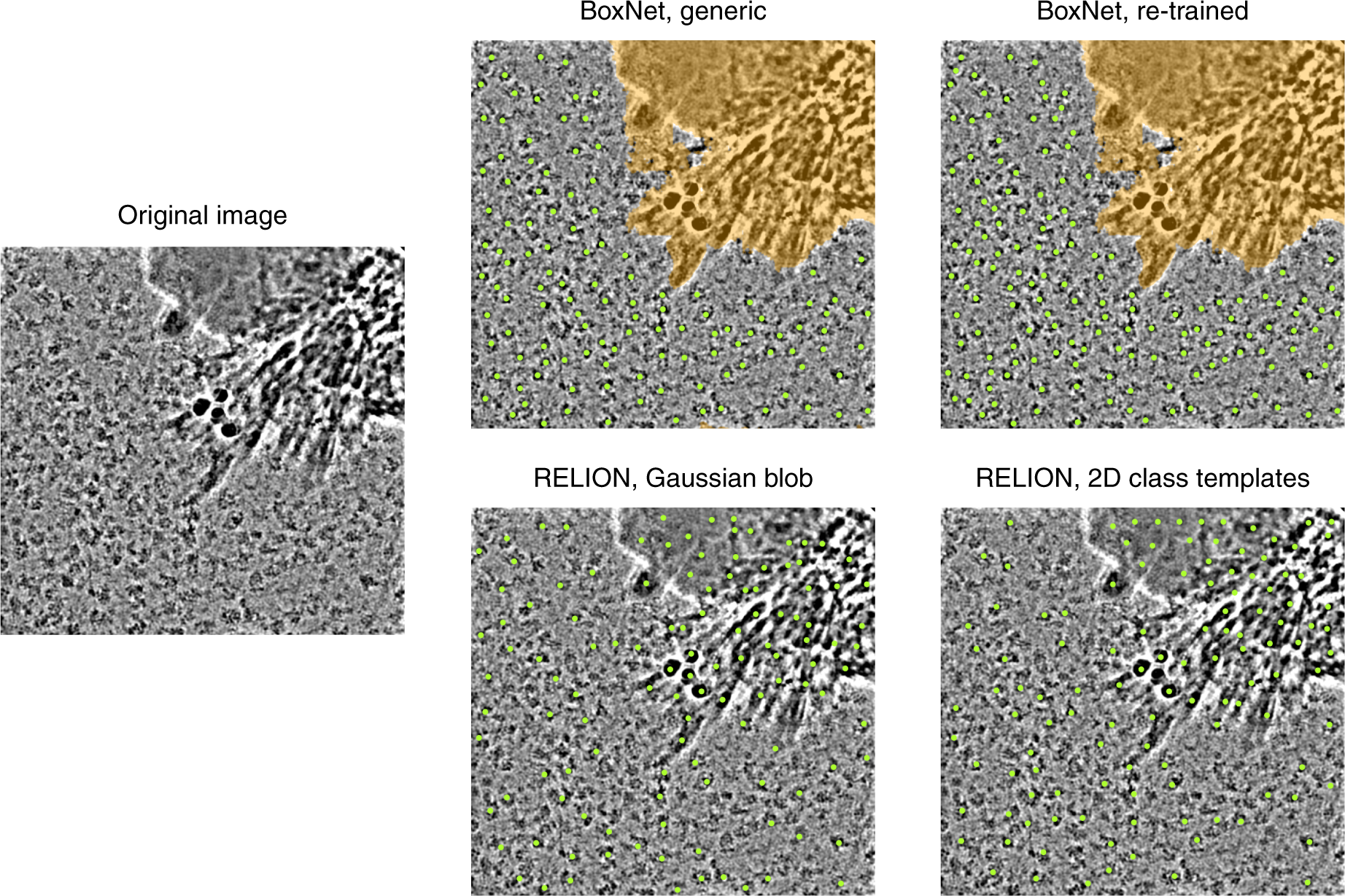
Automated particle picking with *Warp*’s deep learning-based *BoxNet*. Representative example of automated particle picking with *BoxNet* in *Warp* on a micrograph with high-contrast artifacts. Areas masked out automatically by *BoxNet* are colored orange. The generic version of *BoxNet* was never presented with the sample during training. The re-trained version was given 5 micrographs of the same sample, which did not include the one shown. The template-based picking in RELION used 25 class averages derived from 3000 particles, filtered to 20 A. RELION’s results show the 120 highest-scoring positions. For visualization purposes, the micrograph was deconvolved, high-pass filtered and cropped at the borders.

### Retraining of BoxNet

Since the performance of BoxNet can vary between different data, *Warp* offers a retraining interface for BoxNet. Such retraining leads to human-like accuracy in automated particle picking. For retraining, the user can indicate to *Warp* positive and negative examples of BoxNet performance. Using ~1000 examples, retraining of BoxNet typically takes less than 10 minutes, with an estimate of the achieved accuracy provided during the process. After retraining, the user can pick the same micrographs with the re-trained network and select more positive and negative examples for another round of retraining if required. To decrease the need for retraining in the future, *Warp* also provides the option of submitting training data to a central GitHub repository. *De novo* training will be carried out by us periodically with all deposited data, and the resulting updated pre-trained BoxNet offered to the community. The training set is centrally curated and a list of particle species in the current version is available from https://github.com/cramerlab/boxnet. The BoxNet version name will be stored in each micrograph’s metadata to ensure reproducibility of picking results obtained with older versions.

### Online pre-processing during data collection

The design of *Warp* is optimized for processing raw cryo-EM data immediately during data collection. Files written out by the image acquisition software are detected automatically in the specified input folder and added to the list of ‘processable items’ in *Warp*. Each item maintains its metadata in an XML file that includes the exact previous processing settings. Warp continuously performs the processing steps necessary to bring each item into accord with the settings currently specified for the entire folder. During the processing, all results can be immediately inspected. Items can be forcibly included (i. e. exempted from the quality filters) or excluded from downstream processing. The processing must be stopped to change the settings or to retrain the BoxNet model. If changes were made, *Warp* will first reprocess all outdated items. During online processing, *Warp* is able to estimate parameters such as motion, defocus and the resolution limit from micrographs, as well as perform particle picking within less than one minute after the raw data become available. In our experience, high-quality single-particle data of complexes of RNA polymerase II enable the user to obtain detailed 2D classes of particles and 3D reconstructions at better than 5 Å resolution using *Warp* and cryoSPARC within only a few hours after the start of data collection (D. Tegunov, C. Dienemann, and P. Cramer, data not shown).

### Pre-processing tomographic data

*Warp* can also be used to pre-process data from cryo-electron tomographic (cryo-ET) tilt series. *Warp* can reconstruct tomograms from a tilt series and can perform template matching in tomograms with available 3D structures. The (sub)-tomogram reconstruction considers the local CTF, sample distortion and magnification anisotropy (Methods). Additionally, a deconvolved version of the tomograms can be produced using the same interface to help with their visual evaluation. To ensure the CTF model is as accurate as possible, Warp’s CTF fitting procedure goes beyond fitting the tilt images individually. Instead, local patch 2D power spectra from all tilts are fitted simultaneously, with constraints imposed on the inter-tilt angles, and regularizing assumptions made for the progression of phase shift and astigmatism (Methods). The tilt series CTF fitting can also be performed as part of the online processing.

### Template matching

Finding a previously known structure in new data is central to many stages of cryo-EM data processing. The structure must be compared at many different orientations to every position in the new data under the consideration of the CTF. Warp implements template matching only for 3D templates, because matching a set of 2D templates for particle picking is better handled by a neural network such as BoxNet (see above). A template volume can be either provided by the user or automatically downloaded from the EMDB^25^ through the same UI. For template matching, 2D micrographs are subdivided into tiles. Then, normalized cross-correlation is computed between the tiles and 2D projections generated from the 3D template at the specified angular intervals, convolved with the local 2D CTF (Methods). All local correlation peaks with a minimum inter-peak distance corresponding to the template particle diameter, and the corresponding best-scoring template orientations are saved so that the user can later instantly explore different peak thresholds without repeating the costly correlation step. This procedure is also implemented for tomographic volumes, where the local patches are replaced by local sub-volumes, and the local 3D CTF is considered.

### Software implementation

*Warp* is written in the programming languages C#, C++ and CUDA C. The expressiveness of C# and the availability of powerful development tools kept the high-level data management layer brief and maintainable. *Warp*’s rich UI is enabled by the Windows Presentation Foundation (WPF) framework. All performance-critical parts are implemented to run on a GPU. Central data primitives, such as 2D movies and tilt series, and all associated algorithms are wrapped in a stand-alone C# library that we called ‘WarpLib’. The granularity of most of these methods is fine enough to make them useful for applications beyond those implemented in *Warp*. Thus, WarpLib has the potential to speed up the development of future GPU-enabled tools that provide new functionality around the same data. We intend to keep developing *Warp* to enable state-of-the-art, rapid cryo-EM data pre-processing in the future.

### Benchmarking

To test the performance of *Warp*, we reprocessed a published single-particle cryo-EM data set for the influenza hemagglutinin trimer^10^ (Methods) (Fig. 5). We chose this case for benchmarking because the processing of a 150 kDa protein imaged at 40° tilt required a significant amount of manual screening in the original analysis^10^, providing a challenging test case for the *Warp* pipeline. With the original set of 130,000 particles, cryoSPARC reached a similar resolution as that reported in the original analysis (Fig. 5a, b), showing that refinement in cryoSPARC and RELION yields equivalent results for this data set. However, because this particle set and the general particle population both exhibit significant heterogeneity, we draw the comparison between results obtained after subjecting all data to the same 3D classification steps in cryoSPARC (Methods). For the original set, the best class containing 57,346 particles reached a global resolution of 3.9 Å with a B-factor of −200 Å^2^. The same particles, updated with the defocus information from *Warp*, reached a notably higher resolution of 3.5 Å with a B-factor of −170 Å^2^. This suggests that *Warp*’s local CTF model is more accurate than the per-particle CTF fitting in gCTF^2^ used in the original study. *Warp* processing also estimated a narrower range of astigmatism amplitudes (Fig. 5c), in agreement with the assumption of a stable optical system. For the full, completely automated *Warp* pre-processing pipeline, the best class containing 249,495 particles reached a global resolution of 3.2 Å with a B-factor of −170 Å^2^, accompanied by a significantly increased level of detail in the map (Fig. 5a). After classification in cryoSPARC, the best classes contained 45 % and 51 % of all particles in the original EMPIAR-10097 set and Warp’s automatically picked set, respectively, suggesting a similar degree of particle ‘purity’ in the manual and automated approaches. Furthermore, *Warp*’s result demonstrates that tilted data collection can lead to near-atomic resolution – significantly higher than previously established – with minimal effort in the data pre-processing step. Taken together, our results establish *Warp* as a very useful tool for high-performance, automated cryo-EM data pre-processing.

**Figure 5.**
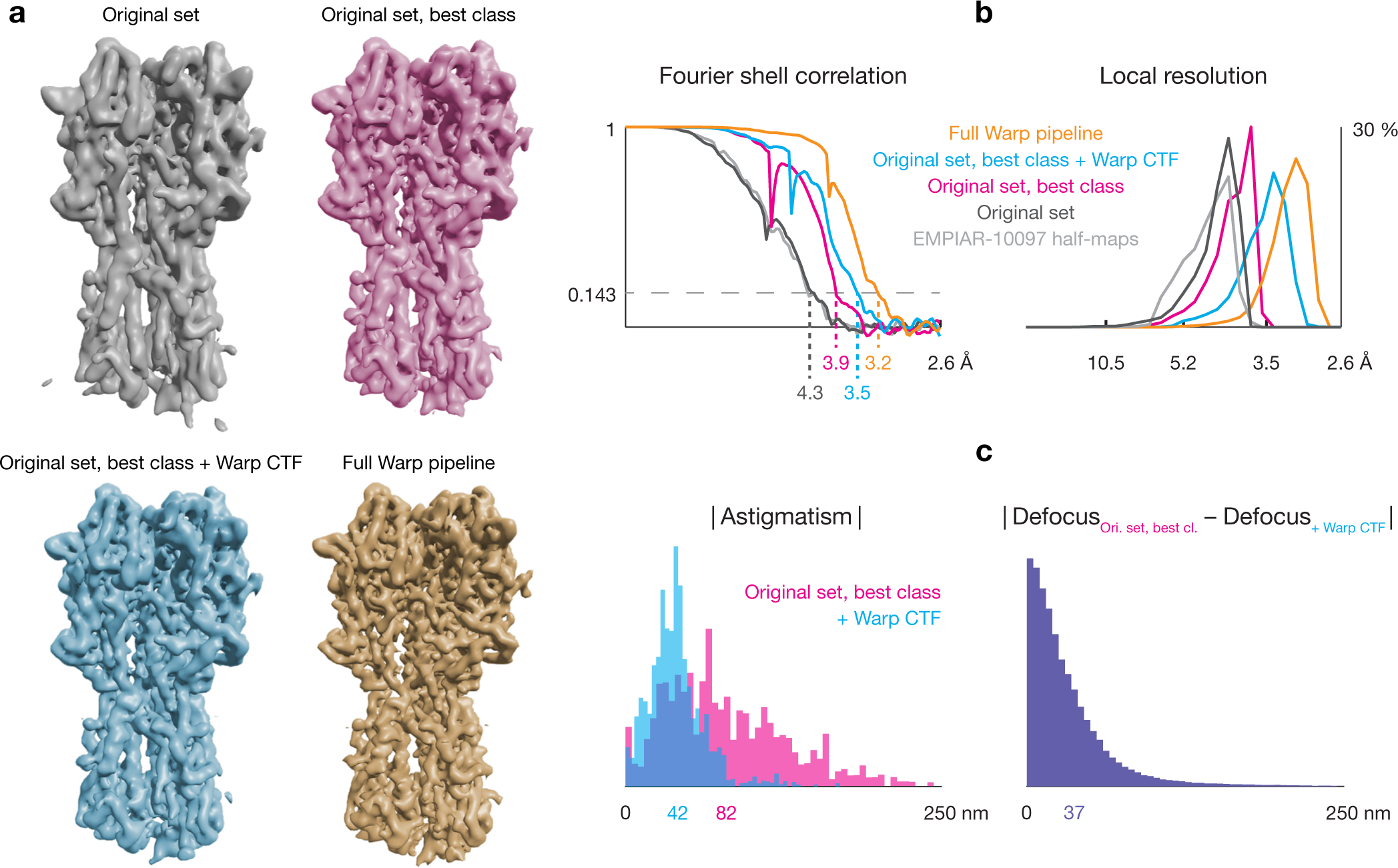
*Warp*’s 2D pipeline improves cryo-EM density for influenza hemagglutinin. As a benchmarking case we used the published EMPIAR-10097 data set containing influenza hemagglutinin trimer particles. ‘Original set’: 130,000 pre-extracted particles from EMPIAR-10097 with their original CTF parameters; ‘Original set, best class’: 57,346 particles from ‘Original set’ after 3D classification in cryoSPARC with their original CTF parameters; ‘Original set, best class + *Warp* CTF’: the same 57 346 particles, but with *Warp*’s CTF estimates; ‘Full Warp pipeline’: 249,495 particles obtained from the raw EMPIAR-10097 data after unsupervised pre-processing in *Warp* and 3D classification in cryoSPARC. (a) Isosurface renderings of the 3D maps generated with cryoSPARC using the respective sets of particles and CTF parameters, filtered to local resolution using the auto-tightened masks generated by cryoSPARC. (b) Global masked FSC plots, and histograms of the local resolution used to filter the maps depicted in (a). ‘EMPIAR-10097 half-maps’ refers to the original half-map volumes deposited in EMPIAR-10097, obtained from the same 130,000 particles as ‘Original set’. (c) Histogram comparison between the original defocus parameters and those estimated by *Warp* for the 130 000 particles from ‘Original set’. The mean value for each metric is specified underneath the horizontal axis in the same color as the corresponding histogram.

## DISCUSSION

Here we present *Warp*, a novel software tool for real-time evaluation, correction and pre-processing of cryo-EM data during data collection. *Warp* bridges between the microscope and available software for 2D classification and 3D reconstruction, taking care of all pre-processing steps. *Warp* also helps the microscope operator to monitor the data quality during acquisition, thus allowing the user to change and optimize the acquisition strategy during ongoing data collection. This enables one to make the best use of microscope time and prevents disappointing results during traditional data processing long after the data collection is concluded. A particularly useful tool included in *Warp* is the deep learning-based algorithm BoxNet that performs reliable, automated particle picking. *Warp* is easy to install, has an intuitive user interface, and will be maintained and improved over the next years.

To further improve cryo–EM data collection efficiency in the future, we envision that an automated feedback loop between the acquisition software and *Warp* would be particularly useful. Such feedback could be beneficial towards the goal to maximize the number of images collected in sample areas with the best data quality, as follows. Areas with low estimated resolution or particle count could be skipped quickly. The stage settling time could be adjusted dynamically based on *Warp*’s motion estimate for the previous image. Furthermore, the time-consuming process of focusing the microscope by tilting the beam could be reserved for large stage shifts only, whereas the defocus in recent images could be obtained from *Warp*’s precise CTF fits and used to adjust the instrument without delaying the data collection. Additional feedback from *Warp* users will also contribute to further development of the software, aimed at serving the rapidly growing cryo-EM community.

## METHODS

### Spline interpolation on multi-dimensional grids

Many methods in Warp are based on a continuous parametrization of 1—3-dimensional spaces. This parameterization is achieved by spline interpolation between points on a coarse, uniform grid, which is computationally efficient. A grid extends over the entirety of each dimension that needs to be modeled. The grid resolution is defined by the number of control points in each dimension and is scaled according to physical constraints (e. g. number of frames or pixels) and available signal. The latter provides regularization to prevent overfitting of sparse data with too many parameters. When a parameter described by the grid is retrieved for a point in space (and time), e. g. for a particle (frame), B-spline interpolation is performed at that point on the grid. To fit a grid’s parameters, in general, a cost function associated with the interpolants at specific positions on the grid is optimized. In the following, we distinguish between 2—3 spatial grid dimensions (X and Y axes in micrographs; X, Y and Z in tomographic volumes), and a temporal dimension as a function of the accumulated electron dose.

### Motion model

Two sources contribute to the observed translational shift between frames in a dose-fractionated image sequence. First, mechanical stage instability leads to rapid shift changes that are uniform within the entire frame. Second, beam-induced motion (BIM) causes slowly changing, local motion. Warp considers the physical properties of both sources in its motion model, using two sets of grids to parametrize the frame shifts and sample deformation. Global motion is described by two grids, X_global_ and Y_global_, with high temporal, and no spatial resolution. The temporal resolution can match the number of frames, or, in case finer dose fractionation is performed to reduce intra-frame motion, the resolution can be lower to regularize a potentially overfitted model. BIM is described by two grids, X_local_ and Y_local_, with a temporal resolution of at most 3, and a spatial resolution of typically 4—5 in both dimensions. The overall shifts required to bring the same object in all frames into a common reference frame are then defined as (X_global_ + X_local_, Y_global_ + Y_local_).

### Global and local motion correction

In the absence of known particle positions and high-resolution reference projections, individual frame patches are aligned to their averages. The movie is subdivided into groups of 512^2^ px patches with a 50 % spatial overlap, masked with a raised cosine. To simplify computation, the images are transformed into Fourier space where complex multiplication replaces translation. For each group, the patches are shifted according to the interpolants at their extraction positions using the current grid values. The average of a group’s shifted patches is then compared to the individual patches to calculate the patch costs as

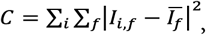

where *i* denotes the frame index, *f* denotes the spatial frequency, *I* is the Fourier transform of a shifted patch frame, and *Ī* is the average of all shifted patch frames. The shifts are obtained by interpolating on the current state of the parameter grids at the patch frame’s position in space and time. The derivative is approximated numerically with the symmetric difference quotient. The overall cost for all grid control points is the sum of all patch costs, and the derivative for each grid control point is the weighted sum of the derivatives of all patches affected by it. The weights for each control point’s derivative can be precomputed by applying a one-pixel shift to the control point and storing all resulting non-zero patch shifts. The cost and derivatives are used by the Limited-memory Broyden-Fletcher-Goldfarb-Shanno (L-BFGS) algorithm^26^ to optimize the values of all control points. The optimization is performed in several steps to improve global convergence. In the first step, the temporal resolution of X_global_ and Y_global_ is set to 3, and increased to the next power of 3 in subsequent steps until the desired temporal resolution is reached.

### Contrast transfer function estimation in micrographs

The CTF analytically describes the convolution applied to the images by the electron-optical system. Estimating its properties with high precision is essential for reversing the effects and obtaining high-resolution reconstructions^27^. Whereas the methodology for measuring defocus and astigmatism from a micrograph’s power spectrum (PS) has been well-established^4, 28^, the recent increase in EM map resolution calls for a more localized approach. Local defocus variation of a seemingly flat sample can exceed 60 nm within a single micrograph, resulting in an out-of-phase CTF for some particles at resolutions beyond 3 Å. Attempts to address this issue by fitting the defocus per-particle have been made^2^, but they require knowledge of particle positions, and lack robustness for all but the largest particle species. Even with a local smoothing approach, per-particle defocus requires high particle density to not lose accuracy compared to a global estimate. On the other hand, strong local irregularities in the specimen surface are almost never observed in tomographic volumes in vitro^17^, suggesting per-particle precision might be unnecessary.

### Estimation of local defocus

To parametrize the defocus, a single grid typically consists of 5×5 spatial control points and 1 temporal control point. Optionally, another single grid with exclusively temporal resolution tracks the development of the phase shift generated by a Volta Phase Plate^29^. Global parameters, such as astigmatism magnitude and angle, are optimized as additional scalars in the model. In practice, the effect of using a more complex geometry or temporal resolution appears negligible. However, the increased signal of future camera hardware might make these options relevant. Like in the frame alignment procedure, groups of patches matching the desired PS size (e. g. 512^2^—1024^2^ px) are extracted with a spatial overlap of 50% from the raw movie data, transformed into Fourier space, and converted to PS by taking their squared amplitudes. If no temporal resolution is desired, each group will be averaged to a single PS to save resources. Similarly, in the absence of spatial resolution, the same frame from all groups will be averaged.

For the initial, exhaustive search, a 1D rotationally averaged PS is calculated from all patches. A B-spline with 3 control points is then fitted through it and subtracted to remove most of the background. The user-defined range of defocus and, optionally, phase shift values is evaluated by matching a simulated 1D CTF. The result with the highest normalized correlation is then used to estimate the 1D PS background and envelope more accurately to consider both in subsequent 2D CTF fitting. The cost function to be minimized is calculated between a 2D PS, and the 2D CTF simulated based on the local defocus at the patch extraction position:

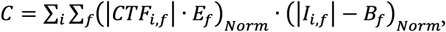

where *i* denotes the frame index (in case a temporal dimension is used), *f* denotes the spatial frequency within the range used for fitting, *CTF* is the 2D contrast transfer function, *I* is the FT of a patch frame, *E* is the envelope of the 1D PS, *B* is the background of the 1D PS, and (…)_*Norm*_ is a normalization operator that brings the value distribution to mean = 0, standard deviation = 1. The defocus and, optionally, phase shift is obtained by interpolating on the current state of the respective parameter grid at the patch frame’s position in space and time. The derivative is approximated numerically with the symmetric difference quotient. The same pre-computed weights strategy as in the frame alignment procedure is employed for the control point derivatives. An L-BFGS algorithm finally optimizes the model for all control point values.

The PS of a tilted plane will usually only show low-resolution Thon rings, regardless of what model was used for the defocus gradient. To provide the user with feedback on whether the more complex defocus model is beneficial, the 2D spectra from all patches, whose parameters are herein referred to as the “source” PS, are rescaled and rotationally averaged to a 1D PS with a single defocus value, referred to as the “target” PS, such that the CTF phases match using the following scaling function:

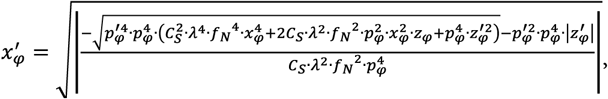

where ‘denotes the “target” PS coordinate system, and its absence denotes the “source” PS coordinate system; *φ* is the sampling angle coordinate, *p* is the anisotropic pixel size, *C_s_* is the spherical aberration, *λ* is the electron wavelength, *f_N_* is the spatial Nyquist frequency, *z* is the anisotropic defocus value, and *x* is the sampling radius coordinate. A similar formulation was provided before^28^ for the special case of isotropic pixel size, and was used to reduce the comparison between CTF and PS to a 1D problem. However, *Warp* performs the fitting in 2D and only uses the rescaling for visualization purposes. If the complex defocus model fits the data better, the recovery of additional highresolution Thon rings can be observed in the 1D average.

### Estimation of resolution

To estimate the useful resolution range, a normalized cross-correlation value between the averaged 1D PS and the simulated 1D CTF curve is calculated within a sliding window. The window size at any given position scales to twice the width between the zero points of the closest CTF peak, but is not allowed to fall below 16 samples. The resolution limit is then reported as the frequency where the cross-correlation falls below 0.3 for the first time. Since the higher number of optimizable parameters allows for some overfitting, it is important that the useful resolution extends beyond the range used for fitting.

### Contrast transfer function estimation in tilt series

The single micrograph CTF estimation procedure with planar sample geometry described in the previous section can be used for tilted 2D data collection. However, full tilt series pose additional challenges for CTF fitting. Mechanical stage instabilities and imperfect eucentric height setting necessitate additional exposures for tracking and focusing^12^ to correct the stage position between individual tilt images. Thus, the defocus cannot be assumed to stay constant, or change smoothly over the course of a tilt series. Each tilt image requires its own defocus value, which can be challenging due to the small amount of signal available. Even at 120 e^−^/Å^2^ for an entire series of 60 images, each tilt only has 2 e^−^/Å^2^ to perform the same estimation as for a 40 e^−^/Å^2^ 2D image, while striving to achieve comparable accuracy.

To improve the fit accuracy, the individual tilt image fits must be subjected to a common set of constraints. As the imaged sample content does not change significantly throughout the tilt series, 1D background and envelope can be derived from the average 1D spectrum of all tilt images. The relative tilt angles and the tilt axis orientation are known to a higher precision than could be derived from fitting a planar geometry *de novo*, and are kept constant throughout the optimization. However, the absolute inclination of the sample plane is unknown. This is especially critical in some of the typical applications of tomography, like the imaging of lamellae prepared through FIB milling because a lamella can be tilted by over 20° relative to the grid. This additional inclination remains constant throughout the tilt series, and is made a single optimizable parameter for all tilt images. Astigmatism and, optionally, phase shift can be kept constant throughout 2D image exposures, but can benefit from a temporally resolved model in a tilt series where the overall exposure is fractionated over a much longer time, e. g. 20—30 min. *Warp* uses 3 control points in the temporal dimension to model these parameters.

With these considerations, the full estimation process is as follows. 2D patches are extracted from all aligned tilt movie averages, as described in the micrograph CTF fitting procedure, and treated in parallel in all subsequent calculations. To provide a better initialization for the per-tilt defocus searches, an estimate for the average defocus of the entire series is obtained by preparing 1D spectra from all patches, and comparing them to simulated CTF curves for the defocus values at the respective positions and tilts, taking into account the fixed relative tilt information and the currently tested average defocus (and phase shift, optionally). This search is performed exhaustively over a range of values specified by the user. The result is used as the starting point of a more complex optimization. Defocus values for all individual tilts, 3 astigmatism magnitude/angle pairs, 3 optional phase shift values, and the two global inclination angles (i. e. the plane normal) are optimized using the L-BFGS algorithm with the derivatives obtained numerically as described in the micrograph CTF fitting section. Upon convergence, the 1D spectra of all patches are rescaled to a common defocus value. This is especially useful for validation in tilt series since the individual images will have very noisy spectra. If the useful resolution range does not extend sufficiently beyond the fitting range, the latter is automatically decreased and the optimization repeated.

In our experience, the direction of the tilt axis is often miscalibrated. Correct handedness in structures obtained from sub-tomogram averaging does not guarantee the tilt angle sign is not flipped. In Warp, a positive rotation around the positive Y image axis is defined to result in an increased underfocus at positions to the positive X side of the tilt axis, i. e. those parts of the sample come physically closer to the electron beam source. The CTF fitting procedure in Warp can detect such mistakes by optionally repeating the optimization with the tilt angles flipped, and notifying the user if the “wrong” set of angles provides a better fit. Such a test can be used to re-calibrate the acquisition software for future data collection.

### Considerations for tomogram reconstruction

Whereas the process of 3D map reconstruction from 2D images of single particles is well established today, full-tomogram reconstruction breaks some of the simplifying assumptions so they must be handled explicitly to obtain better results. In the 2D case, the CTF can be assumed to be the same for all parts of a single particle image, although corrections for a wider range of defocus values in images of large objects have been proposed^30^. In a tomographic tilt series, the highest tilt image can show a defocus spread of 1 um or more. Accounting for such variations in local defocus is necessary for reaching high resolution^31^. Furthermore, each region in the tomographic volume is reconstructed from images with different CTFs, and the zeros and peaks of those CTFs will not overlap in Fourier space. CTF-based weighting of individual projections is commonplace for 2D data^8, 32^, but the algorithms used in tomographic reconstruction do not go beyond CTF phase flipping, giving all spectral components equal weight^31, 33^. This gives spectral components with pure noise (CTF = 0) the same weight as the best available signal (|CTF| = 1) if they overlap. Performing CTF-, dose- and tilt-based weighting later in sub-tomogram averaging has been shown to be beneficial^34^, but it has an even more significant effect when applied at the level of initial tomogram reconstruction. Anisotropic magnification has been described in the past^35^ and is routinely corrected in 2D data. In tomography, the real-space distortion is even more pronounced than in single particle reconstructions because the distances affected by the distortion are more than 1 μm, i. e. the extent of the entire tomogram, leading to positional errors on the order of nanometers in scenarios where the anisotropy does not coincide with the tilt axis.

### Tomogram reconstruction

Warp takes the local defocus and sample distortion, as well as magnification anisotropy into account when reconstructing full or partial tomographic volumes. For a partial reconstruction at any position in the volume, the original 2D images are sampled at the following positions:

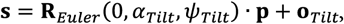

where **R**_*Euler*_ is the rotation matrix for 3 Euler angles following the Xmipp convention^36^, *α* is the stage tilt angle, *ψ* is the in-plane angle of the tilt axis, **p** is the particle position within the tomographic volume, and **o** is the in-plane offset of the tilt axis. The coordinates are centered within the volume and images. The CTF for each 2D image is calculated using a defocus of:

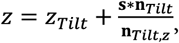

where *Z*_*Tilt*_ is the average defocus estimated for the tilt image, **n**_*Tilt*_ is the sample plane normal, **n**_*Tilt,Z*_ is the *z* component of the normal, and ^*^ denotes the scalar product between two vectors. The reconstruction is performed in Fourier space using a gridding algorithm^8^, with the data weighted by the respective CTF, and the dose- and tilt-dependent heuristic from RELION^34^, but without the final deconvolution step (i. e. the weights are inserted as |CTF|, not as CTF^2^). To obtain a full tomogram, *Warp* reconstructs a uniform grid of small, cubical volumes with an overlap of 50%, and inserts the central 50% into the overall volume to account for artifacts associated with Fourier space reconstruction at the borders of the local volumes. This ensures the corrections can be applied with local precision and remain reasonably continuous between adjacent sub-volumes.

### Export of corrected data

Whereas the *Warp* model for a movie or tilt series describes the non-linear deformation of the entire particle ensemble and its environment, it is unclear whether this deformation gradient stays continuous throughout a single particle, i. e. if the protein structure is subject to the same compression and shearing as the ice around it. Many recent high-resolution maps were reconstructed using particles extracted from dose-weighted averages produced by MotionCor2^1^. The tool assumes the deformation gradient to be continuous in all parts of the image, and will thus deform images of particles and ice in the same way. This will be beneficial if the underlying physical model is indeed continuous. However, it also distorts the CTF locally without passing any knowledge of the distortion to downstream processing tools. In case of a strong local change in the motion direction, this will result in an artifact similar to lens astigmatism.

Warp assumes a continuous deformation field when exporting dose-weighted averages of whole 2D movies, i. e. each pixel will be shifted according to the grid interpolants at that exact position. This has the benefit of uniformly sharper images for visual inspection and particle picking. For particle and sub-tomogram extraction, however, the entire particle image will be shifted uniformly according to the grid interpolants at the particle’s center. This keeps the CTF true to its fitted analytical description, but makes the assumption that the protein is more rigid than the surrounding ice and thus deforms less due to BIM. For whole-tomogram reconstruction, a hybrid approach is pursued: the local volumes are produced using the same procedure as sub-tomogram extraction, but the combined volume is largely continuous depending on how small the local volumes were.

### Particle picking with a residual neural net

In the past years, the recipe for improving the performance of deep learning algorithms has been “deeper networks, more training data”. Outside of cryo-EM, deep ResNet architectures have been demonstrated to enable the training of very deep models by essentially solving the vanishing gradient problem^21^. At the same time, the EMPIAR raw data repository^23^ has accumulated a diverse collection of 2D cryo-EM data sets that can be leveraged for training. *Warp* employs a model with 35 ResNet blocks and 2 conventional convolution layers (Fig. 6) to segment a micrograph into 3 classes: background, particle, and high-contrast artifact (e. g. ethane drops). The input window has a constant size of 256^2^ px. After initial convolution with 32 5×5 kernels the data are processed by a sequence of 5 groups of each 5 ResNet blocks. At the beginning of each but the first group, the data are down-sampled by a factor of 2, while the number of channels is doubled. This enables the recognition of an increasing number of large, higher-order features. After reaching a spatial extent of 16×16 in the contractive part of the network, the data are processed by the expanding part: a sequence of 4 groups of each 2 ResNet blocks. At the beginning of each group, the data are up-sampled by a factor of 2. Each group’s output is concatenated with the output of its mirror counterpart from the contractive part in a U-Net-like fashion^37^. This combines the global context and higher-order features obtained in the contractive part with the higher spatial resolution of the previous layers. After reaching the original extent of 256×256 in the expanding part, the data are processed by the final 2 ResNet blocks and projected onto 3 channels by convolution with 3 1×1 kernels. A pixelwise argmax operation finally retrieves the most likely label at each location in the original image. The model graph and variables are initialized and saved using TensorFlow’s Python API, while all subsequent training and inference are performed through its C API, wrapped in C# classes.

**Figure 6.**
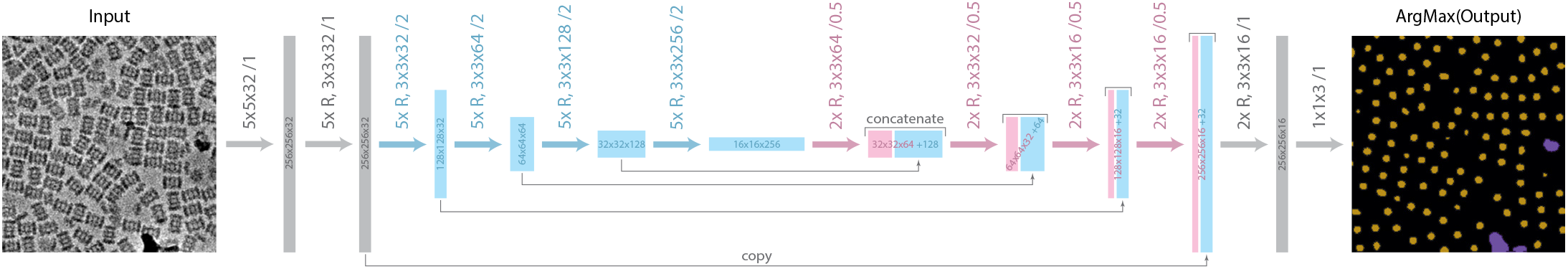
Neural network architecture of *BoxNet*. Rectangles depict the intermediate tensor dimensions. Their width and height are proportional to the number of channels and the spatial extent, respectively. Thick arrows represent convolution operations. Their format is encoded as “(Kx R), LxMxN /O”, where K is the number of consecutive ResNet blocks, or absent in case of a single convolution operation; L and M are the dimensions of the convolution kernel; N is the number of kernels, resulting in N channels in the output; O is the stride length (1 = no change, 2 = downsampling by factor of 2, 0.5 = upsampling by factor of 2 through transposed convolution). The stride parameter is only applied to the first convolution in a chain of ResNet blocks, whereas all subsequent convolutions use stride = 1. The contractive part of the network is colored in cyan, the expanding part in magenta. The final image shows the result of applying a per-pixel ArgMax operator to the result of the last convolution to obtain the spatial distribution of the 3 labels the model is trained to predict: background (black), particle (yellow), artifact (purple).

To segment a micrograph, it is down-scaled to the standard 8 Å/px resolution and divided into 256^2^ px tiles with 64 px overlap. Each tile is extracted, normalized, and presented to the network. All tiles’ softmax and argmax outputs are combined and subjected to a user-defined threshold for the softmax value to remove uncertain picks. Connected components of pixels labeled as “particle” are extracted, and their centroids are used as particle positions. In cases where two particles overlap according to a user-defined diameter, the particle with the bigger connected component is kept. To obtain a mask from the “artifact” label, a threshold of 0.1 is applied to the softmax values, and connected components with less than 20 pixels are removed. The remaining pixels are saved as a binary mask. The final list of particle positions only contains those with a user-defined minimum distance to the masked regions.

### Initial training of BoxNet

10 EMPIAR and 5 in-house data sets (Table S1) each contributed 20–50 micrographs to the training set. Additionally, synthetic data were prepared from 21 PDB models using a modified version of the InSilicoTEM^38^ package, contributing ca. 1600 particles per species. The simulated data contained only one species per micrograph, although more heterogeneous examples might be added in the future. The training set was split 9/1 for training/validation, and trained with the momentum optimizer in TensorFlow 1.5 using a learning rate gradually decreasing from 1E-2 to 1E-5. The normalized data were augmented in each training epoch by extracting the 256^2^ px window at random positions, and applying random rotation, flipping, shearing, and Gaussian noise with a random standard deviation between 0.0 and 0.6. This augmentation was observed to have an excellent regularizing effect, as the final training and validation scores were virtually identical. The training was performed for 800 epochs, using a batch size of 1.

### Retraining of BoxNet

User-supplied positions of positive examples and, optionally, areas of increased and decreased certainty in the micrographs are automatically converted to training data. If requested, the training set is diluted with data from the latest version of the centrally curated set (in the following referred to as ‘old data’) in a 1:1 ratio to prevent possible overfitting of the new data. The retraining regime is identical to initial training, but lasts only 100 epochs. During the retraining, 4 metrics are calculated continuously for every batch: the old network’s accuracy for old and new data, and the retrained network’s accuracy for old and new data. Ideally, the retrained network’s accuracy for new data will improve to approach or even surpass the old network’s accuracy for old data by the end of the retraining process, whereas the accuracy for old data will stay constant.

### Template matching in micrographs and tomograms

2D micrographs are subdivided in tiles with an overlap matching twice the template particle diameter. For each square, 2D projections of the template are prepared at user-defined angular intervals, convolved by the square’s CTF, and normalized to mean = 0, standard deviation = 1 in real space. The square’s FT is multiplied by the conjugate of the projection’s FT, and an IFT yields the cross-correlation scores for all positions within the square. These scores are normalized by the local standard deviation within the square. The scores are compared for all template orientations, and the best one is stored for each pixel within the square. Finally, the result is cropped to exclude a border matching the template particle diameter, and combined with the results from other squares to obtain the correlation scores for the entire micrograph. A local peak search is performed using the template particle diameter as the minimum distance, and all peak positions are stored for further processing. Template matching in tomographic volumes follows the same concept. Instead of square tiles, local cubes are cross-correlated with the template convolved by the local 3D CTF. Optionally, a spectrum whitening of the target micrograph/tomogram can be performed as previously described^39^. This has the benefit of equalizing the spectral noise amplitudes for all spatial frequencies, effectively giving more weight to the higher frequencies and sharpening the correlation peaks.

### Deconvolution

In the absence of a phase plate, the CTF will be dominated by its sine component, i. e. have very little contrast in the lowest spatial frequencies. This creates a high-pass filter effect in the raw data and, due to increasingly noisier higher frequency components, makes it hard to assess the image content visually. On the other hand, a phase plate creating the desired phase shift of π/2 will apply a low-pass filter in defocused images, rendering them blurry. Both scenarios do not affect subsequent alignment and averaging procedures significantly, and the filters will be reversed in the final reconstruction by dividing its 3D FT by the weighted average of all contributing CTFs. This becomes possible because the spectral signal-to-noise ratio (SSNR) is sufficiently high after averaging enough particles with different CTFs. However, even in single images the lowest frequency components often contain enough signal so that boosting them by inverting the CTF will increase the visible low-frequency contrast while maintaining acceptable noise levels. This provides conventional images with a better definition of object boundaries, making their manual selection easier. In defocused phase plate images, this improves sharpness.

To construct a Wiener-like filter, *Warp* makes ad hoc assumptions about the SSNR that can be adjusted by the user. The SSNR is assumed to be a combination of an exponential decay curve and a raised cosine high-pass filter:

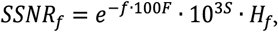

where *f* denotes the spatial frequency, *H* is an optional highpass filter, *F* is the custom fall-off parameter, and *S* is the custom strength parameter. The factors for *F* and *S* are empirically tuned so that the default values of 1 produce good results for typical direct electron detector data, although adjustments might be required in some cases. The SSNR is then used in a Wiener-like filter:

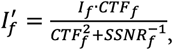

where *I* is the FT of the image, and *CTE* is the 2D contrast transfer function. The shape of the SSNR curve prevents the lowest frequency components from being boosted too much, giving rise to a noisy sample background, and acts as a low-pass filter at the same time to suppress the noisy high frequency components. An example of such a filter and its effect on a 2D micrograph are shown in Fig. 7. In practice, a higher electron dose helps to obtain good low-frequency contrast in conventional images. The commonly used dose of 30–40 e^−^/Å^2^ works well for holey grids with thin ice, while more might be required in the presence of carbon support or thick ice. The deconvolution works especially well in tomograms, where the overall dose often surpasses 100 e^−^/Å^2^.

**Figure 7.**
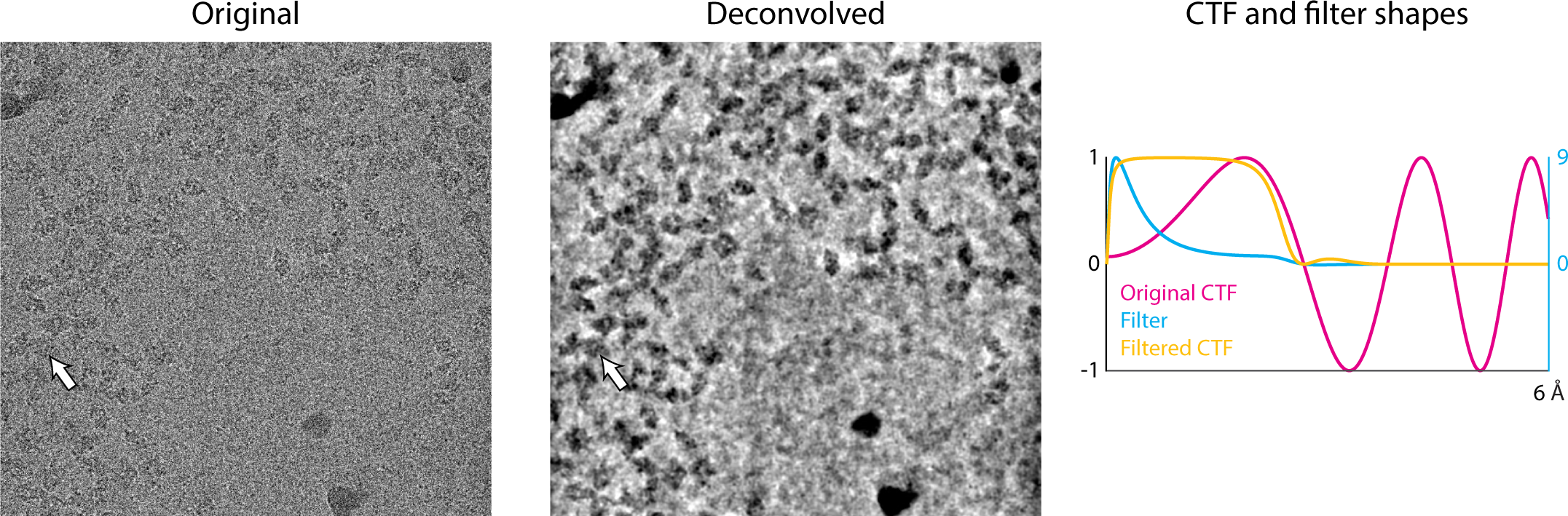
Deconvolution of a low-defocus micrograph. A micrograph from EMPIAR-10061 acquired at 0.9 μm defocus is shown in its original form and after applying the deconvolution filter integrated in Warp. The boundaries of individual 400 kDa proteins can be distinguished more clearly (arrow). The filter largely reverses the effect of the first CTF peak, while also suppressing the lowest and higher frequencies.

### Benchmarking

Raw movie data and pre-extracted particles from EMPIAR 10097 were downloaded. The movies were processed with the full *Warp* pipeline using the following settings: motion correction with a temporal resolution of 40 for the global motion, and 5×5 spatial resolution for the local motion, using the 0.03–0.25 Nyquist range and a B-factor of −400 A^2^; CTF estimation with 6×6 spatial resolution, using the 0.1–0.35 Nyquist range; particle picking with a BoxNet model retrained on particles from 3 micrographs, using the default 0.95 threshold. Quality filters were applied in Warp as follows: defocus between 0.3 and 5.0 µm, resolution better than 8 Å, intra-frame motion of at most 1.5 Å, particle count above 120. Particles were extracted from the micrographs meeting these filters and subjected to processing in cryoSPARC: no 2D classification was performed; *ab initio* refinement was performed with 6 classes and no symmetry; the 6 classes were then refined heterogeneously, with no symmetry imposed; the only class showing the expected Hemagglutinin structure was refined with C3 symmetry. The original particle set from EMPIAR-10097 was subjected to 3 different processing strategies. First, the full set was refined in cryoSPARC with C3 symmetry using the original CTF estimates. Second, the full set was subjected to the same classification and refinement as the particles from Warp, using the original CTF estimates. Third, particles from the Hemagglutinin class obtained in the second processing branch were updated with local CTF estimates from Warp, and refined again with C3 symmetry. Resolution estimates were obtained for all maps using the respective masks automatically generated by cryoSPARC.

## AUTHOR CONTRIBUTIONS

D.T. designed *Warp*’s architecture and all algorithms, and carried out all implementation and application. P.C. provided scientific environment, funding and additional interpretations and implications. D.T. and P.C. wrote the manuscript.

## ACKNOWLEDGEMENTS

We thank members of the Cramer lab for beta-testing early versions of *Warp* and providing feedback on bugs in the soft-ware. We thank C. Bernecky, S. Dodonova, W. Hagen, D. Lyumkis, C. Plaschka, J. Söding and Y.Z. Tan for critical reading of the manuscript. PC was supported by ERC Advanced Grant TRANSREGULON (grant agreement No 693023) of the European Research Council, the Deutsche Forschungsgemeinschaft (SFB 860), and the Volkswagen Foundation.

## DATA AVAILABILITY

The influenza hemagglutinin trimer cryo–EM data used to benchmark the algorithms were downloaded from EMPIAR entry 10097. The 3.2 Å map obtained using the full Warp pipeline was deposited in EMDB as EMD-0025.

## SOFTWARE AVAILABILITY

*Warp*’s binaries, source code and user guide can be downloaded from https://github.com/cramerlab/warp. *BoxNet*’s source code, pre-trained models and training data are available from https://github.com/cramerlab/boxnet.

---

**Table S1.**
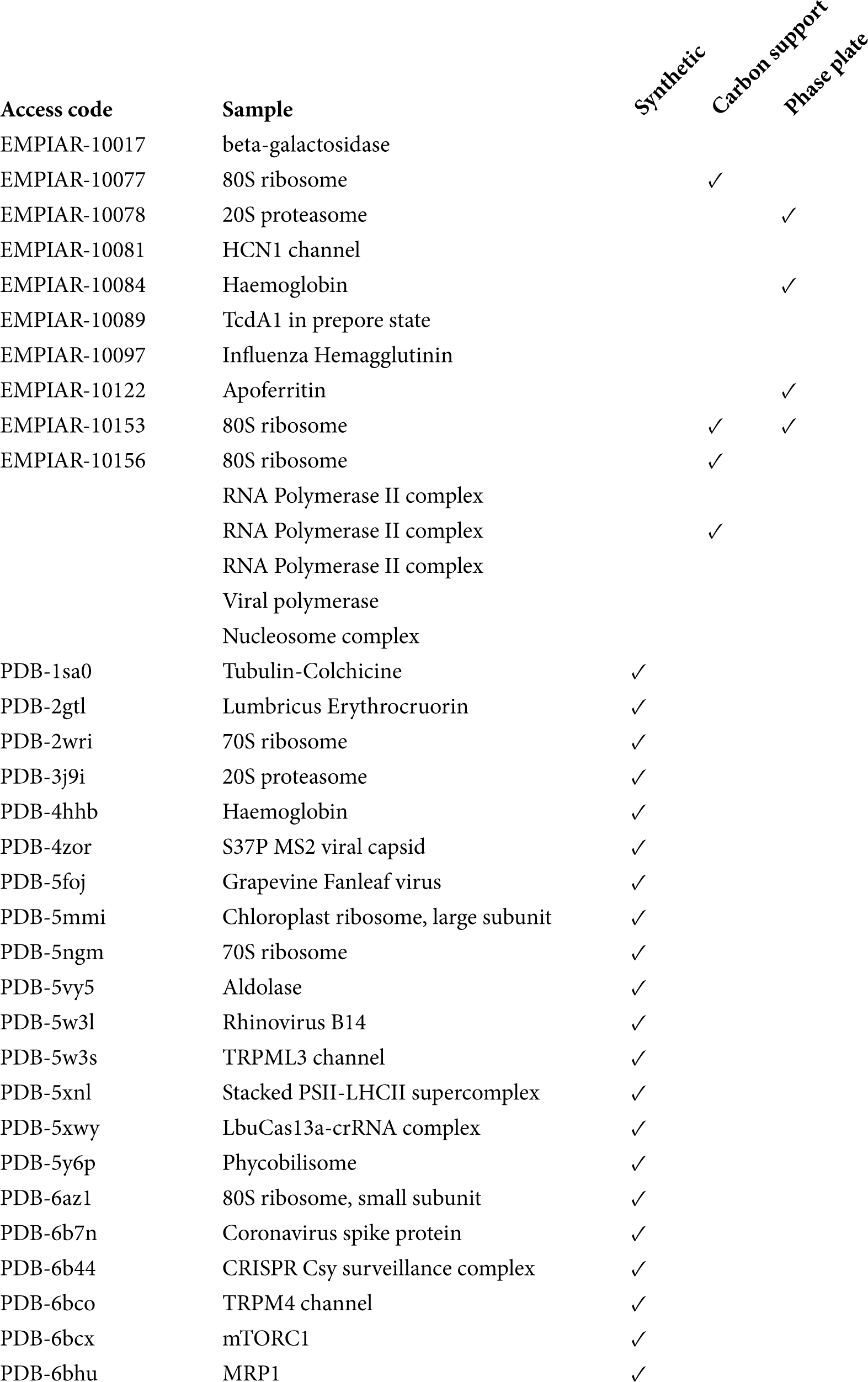
Protein species currently used in the pre-trained version of BoxNet.

